# Genetics of vaccination-related narcolepsy

**DOI:** 10.1101/169623

**Authors:** Hanna M. Ollila, Annika Wennerstrom, Markku Partinen, Emmanuel Mignot, Janna Saarela, Turkka Kirjavainen, Christer Hublin, Logan D. Schneider, Sari-Leena Himanen, Outi Saarenpää-heikkilä, Paivi Saavalainen, Pentti J. Tienari, Outi Vaarala, Markus Perola

**Author notes:** Equal contribution (Hanna M. Ollila, Annika Wennerstrom). Correspondence to: Hanna M. Ollila, 31665 Porter drive, CA 94304 Palo Alto, USA.; Professor Markus Perola, National Institute for Health and Welfare, PL 104, Haartmaninkatu 8, Biomedicum Helsinki, Helsinki 00251, Finland.

## Abstract

Narcolepsy type 1 is a severe hypersomnia affecting 1/3000 individuals. It is caused by a loss of neurons producing hypocretin/orexin in the hypothalamus. In 2009/2010, an immunization campaign directed towards the new pandemic H1N1 Influenza-A strain was launched and increased risk of narcolepsy reported in Northern European countries following vaccination with Pandemrix^®^, an adjuvanted H1N1 vaccine resulting in ~250 vaccination-related cases in Finland alone. Using whole genome sequencing data of 2000 controls, exome sequencing data of 5000 controls and HumanCoreExome chip genotypes of 81 cases with vaccination-related narcolepsy and 2796 controls, we, built a multilocus genetic risk score with established narcolepsy risk variants. We also analyzed, whether novel risk variants would explain vaccine-related narcolepsy. We found that previously discovered risk variants had strong predictive power (accuracy of 73% and P<2.2*10^−16^; and ROC curve AUC 0.88) in vaccine-related narcolepsy cases with only 4.9% of cases being assigned to the low risk category. Our findings indicate genetic predisposition to vaccine-triggered narcolepsy, with the possibility of identifying 95% of people at risk.

---

Narcolepsy is a severe hypersomnia affecting 1/3000 individuals ^1-3^. Narcolepsy type 1 (NT1) is characterized clinically with excessive daytime sleepiness and presence of cataplexy. It is irreversible and caused by a specific loss of hypocretin/orexin producing neurons in the lateral hypothalamus ^4^. Patients with narcolepsy type 2 do not have unambiguous cataplexy and their CSF hypocretin/orexin levels are normal; whereas the hypocretin-deficient NT1 is theorized to be autoimmune-related. In 2009/2010 a vaccination campaign towards pandemic H1N1 Influenza-A was launched. Soon after, an increase in the incidence of narcolepsy was seen in Finland and in Sweden with 6.3/100,000 narcolepsy incidence in children, an 8-fold increase compared to the 0.79/100,000 baseline incidence^5^. The increase was traced back to the use of Pandemrix^®^ for H1N1 immunization ^5-11^. Increased incidence of narcolepsy was subsequently detected in all countries where Pandemrix^®^ was used; whereas countries using other brands did not see increases in narcolepsy incidence ^12,13^. Other environmental risk factors that have been involved in triggering narcolepsy include H1N1 itself and *Streptococcus pyogenes* infections ^14-16^.

The absence of hypocretin neurons in narcolepsy and its association with Influenza-A immunization/infection suggest an autoimmune origin for narcolepsy type 1. This hypothesis is also supported by genetic risk factors that have been identified through Genome Wide Association Studies (GWAS) in narcolepsy. Indeed, nearly all individuals (98%) with narcolepsy type 1 carry the exact same *HLA DQB1*\**06:02* allele ^17-19^ and HLA associations are seen in nearly all autoimmune diseases. Furthermore, variants within the T cell receptor (*TRA* and *TRB*) loci associate with narcolepsy both with and without vaccination as a trigger ^20-22^. TCRs are expressed on T lymphocytes and carry out effector function by interacting with HLA molecules presenting specific epitopes, suggesting a specific antigen presentation pathway for the development of narcolepsy ^21^. Similarly, other known genetic variants (*CTSH*, *P2RY11*, *ZNF365*, *IFNAR1* and *TNFSF4*) interact with the same antigen presentation pathway or predispose to autoimmune diseases ^20-23^.

Finally, many of the genetic effects seen in narcolepsy, such as those for *HLA* and with *TRA*, have unusually high effect sizes with OR>8 and 1.5, respectively. Furthermore, both alleles show association with vaccination-related narcolepsy in Swedish individuals ^24^. Due to the high effect sizes of known variants, we decided to examine, whether individuals with vaccine-related narcolepsy harbored known predisposing risk factors, and whether there were previously undiscovered genetic risk factors specific for vaccine-related narcolepsy cases.

## Materials and Methods

### Patients

The diagnosis of narcolepsy type 1 (ICD-10 code G47.4) was based on criteria of the International Classification of Sleep Disorders (version 3, ICSD-3). In addition, all patients had overnight polysomnography and a positive Multiple Sleep Latency Test (i.e. <8 mins, with 2 or more REM sleep onsets). Absence of hypocretin in cerebrospinal fluid was confirmed in the majority of cases with lumbar puncture (N=58). Informed consent in accordance with governing institutions was obtained from all subjects and reviewed by ethical review boards of Helsinki University, Helsinki-Uusimaa Health District Ethical committee, and Stanford University. The study fully complies with national and international regulations regarding privacy and data protection relevant to medical- and patient-oriented studies. Controls N=2796 were drawn from the Finrisk 2002 and 2007 projects. Coverage of the general population for H1N1 influenza-A Pandemrix^®^ vaccination was 48% ^5,25^.

### Genotyping and Imputation

HLA alleles were imputed using HIBAG and a White specific reference panel ^26^. Posterior probabilities were high: over 95% for each HLA gene. Positivity for DQB1*06:02 was verified directly using a panel of sequence-specific oligonucleotide probes ^27^. Genotyping was performed at the Finnish Genome Center using Human Core Exome chip (Illumina, San Diego, CA, USA) at Finnish Institute for Molecular Medicine (FIMM) technology center genotyping unit or using Axiom EUR (Affymetrix, Santa Clara, CA, USA) at the University of California, San Francisco genotyping Unit. Quality Control (QC) for relatedness and population stratification was performed. Individuals and markers with of > 0.95 missing values and markers with Hardy-Weinberg equilibrium P<0.000001 in controls were removed (N=149 controls and 1 case were removed due to relatedness, population stratification based on PCA, or low genotyping rate). Genotyping rate was 99.8% in the remaining individuals. Finland specific Sequencing Initiative Suomi (SISu) with 2000 whole genome and 5000 exomes was used as a reference for SNP imputation ^28,29^. Pre-phasing was performed using SHAPEIT ^30^ and imputed to whole genome coverage using IMPUTE2 ^31^.

### Statistical analysis

We performed a frequentist association study using SNPTEST ^32^ adjusting for *HLA DQB1*\**06:02* dosage and the first four principal components to adjust for population stratification. Results were compared to analyses without adjustment in addition to analyses stratifying controls to include only *DQB1*\**06:02* positive individuals. To estimate genetic load, we used a multilocus genetic risk score calculation, where the previously reported leading variants at each locus were chosen and weighted by previously reported odds ratio (OR)^20-23^ using PLINK v1.9^33^ and R version 3.2.2 (2015-08-14).

## Results

### Multilocus genetic risk score associates strongly with vaccination-related narcolepsy

We defined significant association at the HLA locus with vaccination-related narcolepsy (P=4.86*10^−44^). Considering presence of *DQB1*\**06:02* in all cases and known increased predisposition with homozygosity ^34^, the remaining analyses were adjusted for the number of *DQB1*\**06:02* alleles.

Altogether seven genomic regions for narcolepsy have been established to date: *TRA*, *TRB*, *IFNAR1*, *P2RY11*, *TNFSF4*, *ZNF365* and *CTSH*, with the T cell receptor having the highest predisposition with an OR > 1.6, across ethnic groups ^20-23^. We examined whether the individual known leading risk variants (**Table 1 and Figure 1**) or the polygenic risk score (Figure 1) from these variants associated with narcolepsy. Some of the individual risk alleles (*TRA*, *CTSH* and *P2RY11*) and the weighted genetic risk score as a whole had substantial predictive power to define vaccination-triggered narcolepsy case vs. control status. The median number of risk alleles was 4 in controls and 6 in cases (Figure 1). There was a strong association of the risk score in the narcolepsy vs. control analysis using the 7 risk SNPs: p=0.0093 and p=0.023 when analyzed against *DQB1*\**06:02* positive controls, and p<2.2*10^−16^ when *DQB1*\**06:02* was included in the model. For predictive power when the cutoff for risk score was set to 0.71 (intercept based on ROC cost-function, optimization to limit false negatives) we could see strong predictive power with 73% accuracy for defining narcolepsy from controls (Figure 1D). The false negative discovery rate was 4.9% with only four cases (4/81) with narcolepsy in the low risk category. On the other hand, 16.6% of controls (495/2796) were assigned in the high risk category, which is substantially clearer separation than with *DQB1*\**06:02* alone, as 28% in the control group carried this allele, yielding high specificity and sensitivity of this genetic risk score model with AUC of 0.88 (Figure 1H). The alleles contributing most to the risk score were *HLA DQB1*\**06:02*, *TRA*, *P2RY11* and *CTSH.* Single SNP level associations were seen with *TRA*; rs1154155 (OR=2.150 [1.524-3.033], P=9.61*10^−05)^ and *P2RY11*; rs1551570 (OR=1.285 [0.914-1.807] P=0.0238) (Figure 1G).

**Table 1.** Replication of individual risk variants for narcolepsy and summarized effects

| Gene | CHR - bp | Reported SNP | Coded allele | P-value | OR |
| --- | --- | --- | --- | --- | --- |
| *TRA* | 14 - 23002684 | rs1154155 | G | 9.61E-05 | 2.150 [1.524-3.033] |
| *TRB* | 7 - 142039520 | rs9648789 | T | 0.192 | 0.810 [0.558-1.176] |
| *CTSH* | 15 - 79234957 | rs34593439 | A | 0.126 | 1.823 [1.096-3.032] |
| *TNFSF4/OX40L/CD252* | 1 - 173131908 | rs7553711 | C | 0.509 | 1.084 [0.765-1.535] |
| *ZNF365* | 10 - 64391375 | rs10995245 | A | 0.443 | 0.943 [0.669-1.329] |
| *P2RY11* | 19 - 10226052 | rs2305795 | A | 0.035 | 1.277 [0.908-1.795] |
| *IL10RB-IFNAR1* | 21 - 34689253 | rs2834188 | G | 0.793 | 0.979 [0.659-1.455] |
| *DQB1* *06:02 | 6 - 32000000 | DQB1*06:02 | *Present* | $2 \times 10^{-54}$ | INF |

odds ratio (OR), base pair (bp), CHR (chromosome)

**Figure 1.**
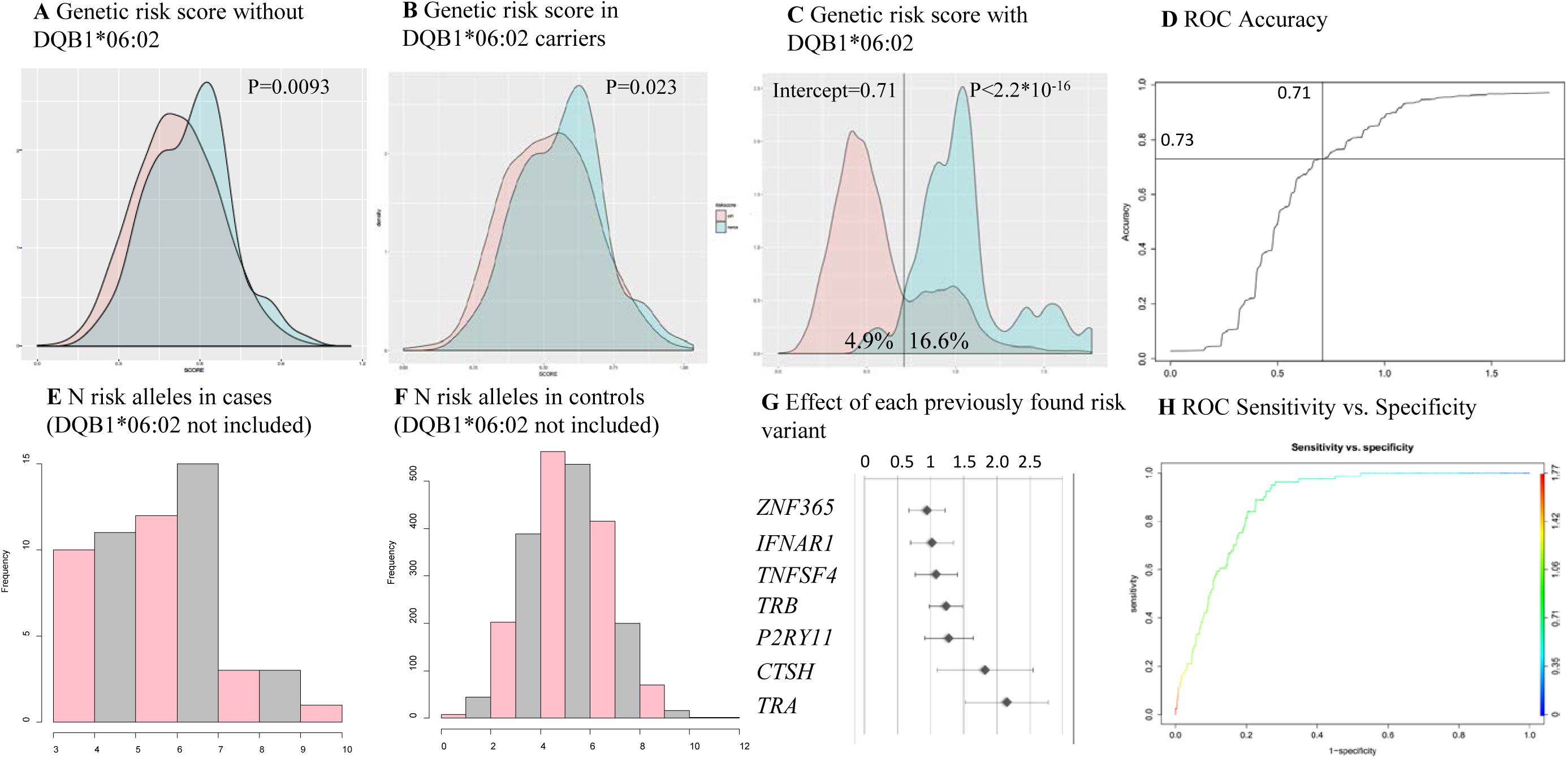
**(A)** Genetic risk score without *HLA DQB1*\**06:02* associates with narcolepsy case control status. **(B)** The separation is seen also when stratifying the controls to be *DQB1*\**06:02* positive **(C)** By including both previously discovered genetic risk factors and HLA DQB1*06:02, strong predictive value is seen with only 4/81 individuals with narcolepsy (4.9%) false negative and 495/2796 controls (16.6%) false positive; 28% of controls and 100% narcolepsy cases were positive for the *DQB1*\**06:02* marker. **(D)** The categorization accuracy of the genetic risk score was 73%. The median number of risk alleles **(E)** in cases was 6 vs. **(F)** 4 in the controls. **(G)** Association of individual risk variants from previous GWAS studies in vaccination-related narcolepsy. **(H)** ROC curve analysis confirms that the overall score had high sensitivity and specificity with AUC 0.88.

### Common variants affect disease risk in vaccination-related narcolepsy

We finally examined, whether distinct genetic variants would underlie vaccination-related narcolepsy. In the full GWAS, three loci close to genome-wide significance level appeared: notably, a highly protective intronic variant in *IL12RB1*, which was present in 1.4% of cases vs. 11% of controls (rs17885060, OR=0.106 [0.026-0.43], P=2.93*10^−7^, Figure S1, Table S1). In addition, a SNP at *ZXDC*, a gene required for HLA class II transcription, was among the top variants (rs11715293 P=7.06*10^−7^, OR=3.33[2.14-5.20], as were variants at *RBM47* (OR=2.74[1.92-3.91], P=1.82*10^−7^) and *SORBS2* (OR=2.14[1.50-3.07], P=3.30*10^−7^). Although not meeting criteria for GWAS significance, the fact that 3 of the 4 associated variants are in known immune regulatory genes was notable. Further, analysis of the biological consequences of these variants defined that the rs11715293 variant on *ZXDC* is a STAT binding site and the risk allele was an eQTL for increased *ZXDC* expression in whole blood (Z= 8.23, P=1.84*10^−16^)^35,36^. For *IL12RB1*, the most likely causative variant is the SNP with second-strongest association at the *IL12RB1* locus rs3761041 (LD r^2^=1 with rs17885060). This variant is in interferon binding site, and an enhancer and H4K3me1 methylation site in the thymus and on T-cells, based on Encode methylation data and HaploReg ^36^.

## Discussion

In this study we discovered that previously reported narcolepsy risk variants could distinguish controls from cases of narcolepsy that have developed the disease following pandemic H1N1 vaccination, specifically with the Pandemrix^®^ brand vaccine. This was established in a new, independent sample of cases that had not been tested for these variants. Everyone with narcolepsy was *HLA-DQB1*\**06:02* positive ^17^. However, the addition of the 7 other known risk variants substantially improved separation between cases and controls. For comparison, the *HLA*-*DQB1*\**06:02* allele is present in 28% of controls while only 17% of controls have a predictive score higher than the optimal threshold for predicting narcolepsy. Furthermore, using this clinically relevant threshold, fewer than 4.9% of narcolepsy cases were estimated to be within the control group resulting in 73% overall accuracy and high sensitivity and specificity. This is important to ensure potential cases of vaccine-inducible narcolepsy are not missed.

The resolution of separation with a genetic risk score is surprising and substantially large. In current medical practice *HLA DQB1*\**06:02* is often used as a tool to help with the diagnosis of narcolepsy type 1. Using the genetic risk score that we built may provide higher resolution in defining those with genetic load. From a clinical standpoint, this is highly valuable, as a polygenic risk score may help us identify individuals that are in risk for developing this permanent and devastating disease.

The fact that large genetic effects were discovered with common variants, together with the fact most (75%) twins are discordant for narcolepsy^37^, suggests that a combination of specific environmental triggers for narcolepsy must occur at the right time in combination with genetic predisposition. Given that only 1 in 16,000 vaccinated children ever developed narcolepsy (versus ~0.71 for 1000,000 at baseline) in Finland^5^, the environmental exposure alone appears insufficient to trigger the elevated narcolepsy incidence noted during the Pandemrix^®^ vaccination campaign. Even in subjects with a genetic threshold above our cut off threshold, only 4 of the 10,000 of highly predisposed subjects would develop narcolepsy following Pandemrix^®^. Clearly, although Pandemrix^®^ was a strong trigger in this population, most predisposed patients never developed the disease, possibly explaining why risk varies substantially following vaccination or H1N1 infections in various countries ^7-10,12,13,24,25^. The other, environmental, immunological, and genetic factors that contribute to the risk of vaccine-related narcolepsy remain unknown.

Of special interest was the fact that two of the individual risk variants (*TRA* and *P2RY11*) independently associated with vaccination-related narcolepsy ^22,23^. Using all previously associated markers, we built a risk score estimating total genetic load and this score had strong independent effects in vaccination-related narcolepsy cases. Importantly, the effect size for *TRA* was higher in vaccination-related narcolepsy, with OR > 2.5, compared to OR = 1.6 in earlier studies ^20-22^, which did not focus on vaccine-related narcolepsy. The association with *TRA* was also observed in a smaller post-vaccination study reported in Sweden, suggesting differential genetic predisposition for various, known, narcolepsy-associated SNPs ^24^.

Additionally, this study provides insights that may be specific to the pathogenesis of vaccination-related narcolepsy. First, when examining the association of variants that may be potentially specific for vaccination related narcolepsy, we found protective variants in *IL12RB1* that were nearly absent in individuals with narcolepsy, with allele frequency of 1.4% in cases but over 11% of controls. This is likely significant as prior studies have shown and replicated the findings of increased measles and pertussis toxin vaccine responses in subjects carrying specific *IL12RB1* receptor and *IL12* polymorphisms ^38-40^. Furthermore, we found 3 additional loci nearing GWAS significance (*RBM47*, *SORBS2* and *ZXDC*), one of which, a low frequency variant in transcription factor *ZXDC* is functionally relevant to narcolepsy pathogenesis, as it is known to increase the expression of HLA class II alleles, such as *DQB1*\**06:02* ^41^. This variant, rs11715293, was highly enriched in cases (OR>3). The biological consequences of both the *IL12RB1* and *ZXDC* variants are interesting and likely involve increased antigen presentation to CD4^+^ T cells in vaccine-related cases, similar to *TRA* and *P2YR11.* However, although functional evidence suggests that *ZXDC* may affect the predisposition to vaccination-related narcolepsy this finding, as well as those of *RBM47* and *SORBS2*, would need to be confirmed through replication in a larger sample set of vaccination-related narcolepsy cases in comparison to narcolepsy not related to vaccination.

## Acknowledgements and Funding

We wish to thank Anne Huutoniemi and Minttu Sauramo for sample and data collection and coordination. This work has been supported by The EU FP7 under grant agreements nr. 313010 (BBMRI-LPC), no. 305280 (MIMOmics), and HZ2020 633589 (Ageing with Elegans), The Finnish Academy grant no. 269517 The Yrjö Jahnsson Foundation (for MP). Finnish Cultural Foundation, Sigrid Juselius Foundation, Orion Research Foundation, Jalmari and Rauha Ahokas Foundation and Academy of Finland (HMO). NIHP50-NS023724-19A1 (EM). NIHT32 HL110952 05 (LDS).

